# Core bacteria associated with hyphosphere of *Fusarium oxysporum* f. sp. *niveum* over spatial and temporal differences

**DOI:** 10.1101/2023.08.11.552976

**Authors:** Vanessa E. Thomas, Sanjay Antony-Babu

## Abstract

**Background:** Bacteria and fungi co-inhabit the soil microbiome in dynamic interactions. In the rhizosphere, fungi and bacteria have been studied to synergistically colonize soil as beneficial or as antagonists to form a pathobiome. These variations of soil bacterial community from pathogen and nonpathogen form of FOSC have been researched, however the bacterial community within the hyphosphere has yet to be studied thoroughly for direct pathogen interkingdom interactions. This study used 16S rRNA gene sequencing and a to decipher the bacteriome diversity associated with the hyphosphere of three isolates of *Fusarium oxysporum* f. sp. *niveum* race 2 (FON2) with temporal and spatial differences.

**Results:** Our results show a core microbiome that is shared among the three isolates regardless of the differences of spatial and temporal differences. The core hyphosphere community visualized as a ternary plot was made up 15 OTUs which were associated with all three FON2. Although a few operational taxonomic units (OTUs) were significantly correlated with a particular isolate of FON2, reported in the LDA (p<0.05), these OTUs were still present as part of the core in all isolates. Co-occurrence analysis and correlation plot identified a negative correlation among most of the microbiota which may indicate a positive correlation to the FON2 that is not tested.

**Conclusions:** The study indicates a core microbiota associated with FON2 regardless of the isolate’s temporal and spatial differences. Through our results we provide insights into the microbe-microbe dynamic of the pathogen’s success and its ability to recruit a core pathobiome. Our research promotes the concept of pathogens not being lone invaders but recruits from the established host microbiome to form a pathobiome.

## Background

Hyphosphere represents a major hotspot of interkingdom interactions between fungi and bacteria. The hyphosphere environment is a zone of influence in the soil that is immediately around a fungal hypha, where exometabolic chemistry is highly altered by the host fungi and hyphal-associated bacteria [1, 2]. Within the agricultural systems, the hyphosphere represent a major micro niche [3]. Bacterial physical association with the fungal hypha can be varible, including: freely motile and moving along the hyphae, attached as single cells, within the hyphae as endohyphal bacteria, or forming biofilms along the hyphae [3, 4]. The planktonic bacterial movement on the hyphae, termed the hyphal highway, has been observed in several soil fungi. For example, the hyphosphere of soil fungi *Lyophyllum* sp allows several individual bacteria to move along the hyphae and form biofilms [5]. Well-studied bacterial fungal interactions (BFI) are among “helper bacteria” and ectomycorrhizal fungi, which synergistically increase soil and plant health [6, 7]. These beneficial bacteria can antagonize phytopathogenic fungi, such as *Pseudomonas fluorescens* (Within a separate study a different isolate of *Pseudomonas fluorescens* was previously cited as a helper bacterium of ectomycorrhizal fungus *Laccaria bicolor*), which reduces a range of fungal pathogens through the production of antimicrobial polyketides [8, 9].

Similar to bacterial and fungi forming plant-beneficial relationships, bacteria and fungi have been studied to form phytopathogenic alliances to infect their host [10]. Within the hyphosphere bacterial alliances may be a strategy to assist in pathogenicity; evidence of such a with *Burkhholderia* endosymbiont of rice seedling blight (Rhizopus microspores) by producing phytotoxins allowing the fungi pathogen to infect the plant host [11]. Other synergistic alliances include Wheat pathogen *Septoria tritici* has been studied to recruit *Erwinia carotovora atroseptica* to increase symptoms within infected hosts [12]. Further studies within this field include *Enterobacter sp.* with soil-borne phytopathogen *Rhizoctonia solani* and *Fusarium oxysporum* f. sp. *lactucae* with its assemblage of ectosymbiotic bacteria [13].

In the study presented here, we set-out to investigate the hyphosphere bacterial diversity of Fusarium wilt pathogen of watermelon. A wide range of crop plants are known to be affected by Fusarium wilt. The disease is caused by *Fusarium oxysporum* species complex (FOSC), one of the most devastating plant pathogens that is, ranked 5^th^ among the top 10 plant pathogenic fungi [14]. *F. oxysporum* f. sp. *niveum (FON)* is host specific to watermelon. In states such as Florida, Georgia, and Texas where watermelon cultivation is prevalent, farmers can experience 30-80% or at times total yield loss from this soil pathogen [15–17]. As a soil-borne pathogen, it is reasonable to assume that FON2 will interact with diverse bacteria, with some involved in assisting in its pathogenicity. Towards evaluating the hyphosphere bacteria acting as a FON’s pathobiome, as an initial step, we investigated whether the hyphosphere bacterial community of the fungal inoculum from the infected tissues. As infected tissues as inoculum within plant residues, we envisaged that the bacterial associates in this fungal inoculum would best represent the pathobiome. We will specifically answer the following questions: 1) does FON maintain a hyphosphere bacterial community? 2) if so, can the fungi maintain the bacterial diversity consistently through several subcultured generations? 3) do different isolates have a central core bacterial community? Finally, 4) is there evidence of strain specific interactions between fungi and bacteria? In the study presented here we define the hyphosphere bacterial assemblage of watermelon Fusarium wilt in three different FON isolates from different counties in Texas from 2002-2020 (COM 2020, W2 2018, and HAR 2002). This study sets the stage to study hyphosphere bacteria of soil-borne fungi as potential pathobiome of the fungal pathogen. Methods

### Seed and Soil Sterilization

Watermelon seeds (Super Gold) acquired from Dr. Thomas Isakeit (Texas A&M AgriLife Extension) were washed with Tween 20® (0.002%), rinsed with 10% bleach, surface sterilized with 70% ethanol and rinsed with sterile water; with each done separately for 2 minutes. After sterilization, watermelon seeds were placed on sterile 2% gelrite in Petri dishes. Aliquots of 100ul of ¼ Hoagland’s solution were added to each seed [18]. Seeds were incubated in a dark box for up to 7 days at room temperature. The soil mixture with 1 part play sand, 1-part commercial soil (Jolly Gardener Pro-Line C/25 Growing Mix) was autoclaved (gravity cycle 60 min) once per day for three consecutive days. The sterile soils were moistened with 5ml of sterile ¼ Hoagland’s solution into which the germinated seeds were planted. Plants were kept in a growth chamber with light intervals of 10 hours of dark and 14-hour light.

Inoculation of watermelon plants

#### FON2 inoculum preparation

We used 3 FON2 isolates (COM, HAR, and W2) from our earlier study [19]. Briefly, COM was isolated in 2020, W2 in 2018, and HAR in 2002. All Isolates were initially isolated from symptomatic watermelon plants. In this study, the FON2 inoculum was prepared by culturing the fungal strains in tryptic soy broth (TSB) on an orbital shaker for 7 days at room temperature.

#### Inoculation into sand-oatmeal matrix

A growth matrix consisting of soil and oatmeal was prepared. This matrix included an equal mass of oatmeal (The Quaker Oat Bran™) and play sand. This mixture was dispensed as 10 grams per glass Petri plates and sterilized by autoclave. Following this, 200ul of FON2 inoculum and 5ml of 1x phosphate buffer solution (PBS) were dispensed into sand-oatmeal matrix and incubated at 27**°**C for up to 7 days. To inoculate the soil, a part of the soil near the rhizosphere was removed without disturbing or touching the roots. This soil was mixed with the sand-oatmeal fungal mix, and reintroduced to the initial site from where the soil was removed.

### Tissue sampling, and generation culturing from the hyphal tip

After the 10-day post-inoculation, stem slices (>3mm thick) from the infected and non-infected plants were removed from above the soil line. The tissue samples were rinsed with 10% bleach, surface sterilized with 70% ethanol and washed with sterile water, each step done separately for 2 minutes before aseptically transferred to the center of a half-strength potato dextrose agar (½ PDA). The plates were incubated upright in an incubator at 27**°**C for seven days. These plates were labeled “*Generation 1*” cultures. On day seven, when hyphae had grown from diseased tissue, sterilized toothpicks were used to carefully collect the hyphal tips at the edge of the colony from the surface layer of the agar media at 4 quadrants to represent 4 replicates per plate. Hyphal tip microbiota collected from sterilized toothpicks were suspended in molecular-grade water (1 mL). Then, 20ul of the FON2 water suspension was dispensed to the middle of ½ PDA plate labeled as *Generation 2*. Finally, 500ul of the water and FON2 solution was mixed with 500ul of 60% glycerol and stored at -80**°**C for preservation, and the remaining amount was stored as our DNA material for amplicon-sequencing. Of the four *Generation two* plates 3 were resampled at 4 quadrants as earlier and labeled as *Generation three* plates. This resulted in 13 microbiota suspensions that were later used for microbial community analyses.

### Microscopy

Samples were viewed with the Olympus BX60 compound microscope using 40×/0.65 Ph2 objective and images were taken using the Olympus camera model LC30 (Olympus Soft Imaging Solutions GMBH, Munster, Germany). Samples imaged were of FON2 COM isolate grown on 0.5X PDA, under the microscope images of the hypal tip were taken to view bacterial hyphal interactions.

### DNA extraction and SSU rRNA library preparation

Aliquots of 200ul of 13 suspensions were used as DNA samples which were placed at -18°C for freeze-thaw lysis-based approach. DNA quantification was assessed in a SpectraMax ® QuickDrop™ Micro-volume spectrophotometer (Molecular Devices, USA). DNA samples were then quantified with Qubit® 2.0 Fluorometer (Invitrogen, Life technologies, USA) with AccuGreen™ High Sensitivity dsDNA Quantitation Kit (Biotium, USA). The presence of fungal and bacteria DNA within the samples were assessed by PCR using established universal primers: ITS1 (5’-TCC GTA GGT GAA CCT TGC GG-3’) and ITS4 (5’-TCC TCC GCT TAT TGA TAT GC-3’) for fungal DNA, and 16S rRNA gene amplification using 27F (5’-AGA GTT TGA TCM TGG CTC AG-3’) and 1492R (5’-GGT TAC CTT GTT ACG ACT T-3’) primers for bacteria [20, 21]. A nested amplification approach was used to prepare amplicon sequencing libraries for 16S rRNA. The first step of nested PCR utilized primers 27F and 1492R, samples with 16S rRNA gene amplification and were visually confirmed after electrophoresis on 1% agarose gel containing GelGreen (Biotium, USA). The second PCR reaction consisted of 16S rRNA gene primers without overhang 341F (5’-CCT ACG GGN GGC WGC AG-3’) and 785R (5’-GAC TAC HVG GGT ATC TAA TCC-3’) [22]. PCR conditions followed the KAPA HIFI Hotstart Readymix PCR kit protocol (KAPA Biosystems). Final nested PCR products were sent to Novogene (Novogene Corporation Inc., Sacramento, CA, USA) for 16S rRNA gene sequencing with Illumina NovaSeq PE250 platform (Illumina, San Diego, CA).

### Sequence analysis

Metabarcoding sequence analysis were processed in Mothur v.1.48.0 [23] following the MiSeq SOP pipeline with modifications [24]. The modifications introduced were based on our initial assessment with sequences obtained from the 13 hyphosphere of one of the tests FON2 strains, COM. Based on the initial rarefaction curves (Fig. S1) we determined that sequence depth of 50,000 reads per sample would suffice for hyphosphere metabarcoding. Additionally, in the initial test the align.seqs step resulted in loss of several reads which was rectified when the “flip” option was set to “flip=T” (to reverse complement the reads that did not align in the forward direction). Based on these observations, a modified pipeline incorporating these two changes was applied to the final analyses of the 39 samples (13 samples per FON2 hyphosphere). The OTUs thus obtained were classified with SILVA v132 bacterial reference trained with our v3-v4 primers.

### Statistical Analysis

Mothur OTU results were analyzed and visualized using R version 4.2.1 (RStudio, Boston, MA, USA). Alpha diversity indices were calculated within Mothur to obtain the richness and evenness measures within our samples, including Shannon, Chao, and Simpson evenness. Tabulated alpha diversity output from Mothur was compared among isolates with ANOVA and Tukey HSD test and visualized using ggplot2, stats, and multicompview packages [25, 26]. Relative abundance stats obtained from Mothur were utilized in co-occurrence network analysis. Three different networks were generated for each of the individual’s hyphosphere populations. The networks were generated as Spearman correlations (coefficients >0.7) within the Microeco package and visualized with Gephi 0.10.1 [27, 28]. FASTA output from “get.oturep” command to get list of OTU and representative sequences were utilized for Maximum-likelihood phylogenetic trees and calculated within molecular evolutionary genetics analysis program (MEGA11) and utilized within the interactive tree of life program for tree visualization of the core microbiome among isolates (iTOL v6) [29, 30]. Weighted and unweighted UniFrac were visualized with r packages phyloseq, tidyverse and ggplot packages with maximum-likelihood phylogenetic trees made from MEGA11 program. Linear Discriminant Analysis (LDA) with logarithmic score significant value of 0.05 and LDA cut off of 3 was performed at OTU level to identify taxa that are significantly unique to our isolates hyphosphere; results were visualized with microbiomeMarker package [31].

Core microbiota associated with the hyphal tip was determined by OTUs that occurred in at least 3 samples of the 39 iterations across all three isolates samples. Once the core microbiome was identified, the OTUs of interest underwent arcsine transformation and presented as a ternary plot to visualize the distribution of the microbial communities among our three isolates through the ggplot2 extension ggtern package [32]. Heatmaps were generated from the core microbiota to visualize the relative abundances of the OTUs among the test fungi using the phyloseq package [33]. An OTU was deemed to be core if it was observed in three or samples within the 13 samples. These were visualized as correlation plots to distinguish positive and negative interactions within each FON2 isolates core microbiota; this was generated with corrplot package [34]. Functional prediction of bacterial communities within individual hyphospheres was performed using PICRUST2 based on the Kyoto Encyclopedia of Genes and Genomes (KEGG) database. The t-test was utilized to determine significant differences in functional potential of the hyphosphere microbiota among the three test fungal isolates with RStatix R package [35–37].

## Results

Sequencing of the 16S rRNA fragment from the V3-V4 region yielded 6,272,775 raw reads from our 39 samples. After sub-sampling 50,000 reads and processing through the modified Mothur pipeline resulted in 13,619 reads classified as 1,201 OTUs. The hyphosphere bacterial composition was classified as 5 phyla and 21 families, *Proteobacteria* was by far the dominant phyla (97%) followed by unclassified bacteria (1.16%). At the family level, *Oxalobacteraceae* (39.57%), *Burkholderiaceae* (27.72%), and *Comamonadaceae* (24.47%) were the majority of OTUs identified from all samples.

Alpha diversity indices Chao (Figure S3), Shannon (Figure 1A), and Simpson evenness (Figure S4) were compared to the overall diversity among the three FON2 isolates hyphosphere. Within Shannon diversity analysis, there is a significant difference among the diversity of the FON2 isolates; when comparing COM and W2 they are significantly different (P < 0.05) whereas COM compared to HAR and HAR to W2 were not significantly different (Figure 1E). LDA was conducted at the OTU level to identify possible unique microbial assemblies among the FON2 isolates (Figure 1F). Among the three FON2 isolates Otu002 (*Comamonadaceae*) was highly correlated to COM, while W2 significantly correlated to Otu003(*Burkholderiaceae*). Furthermore, Otu0021 (*Oxalobacteraceae*), Otu0007 (*Oxalobacteraceae*), and Otu001 (*Oxalobacteraceae*) were significantly correlated with HAR. Unweighted UniFrac analysis of the hyphosphere communities was not significantly different regardless of FON2 hosts or generations. Therefore, the community structure from the FON2 isolates is not dissimilar even though the fungal host background has temporal and spatial differences (Figure 1C).

**Fig. 1.**
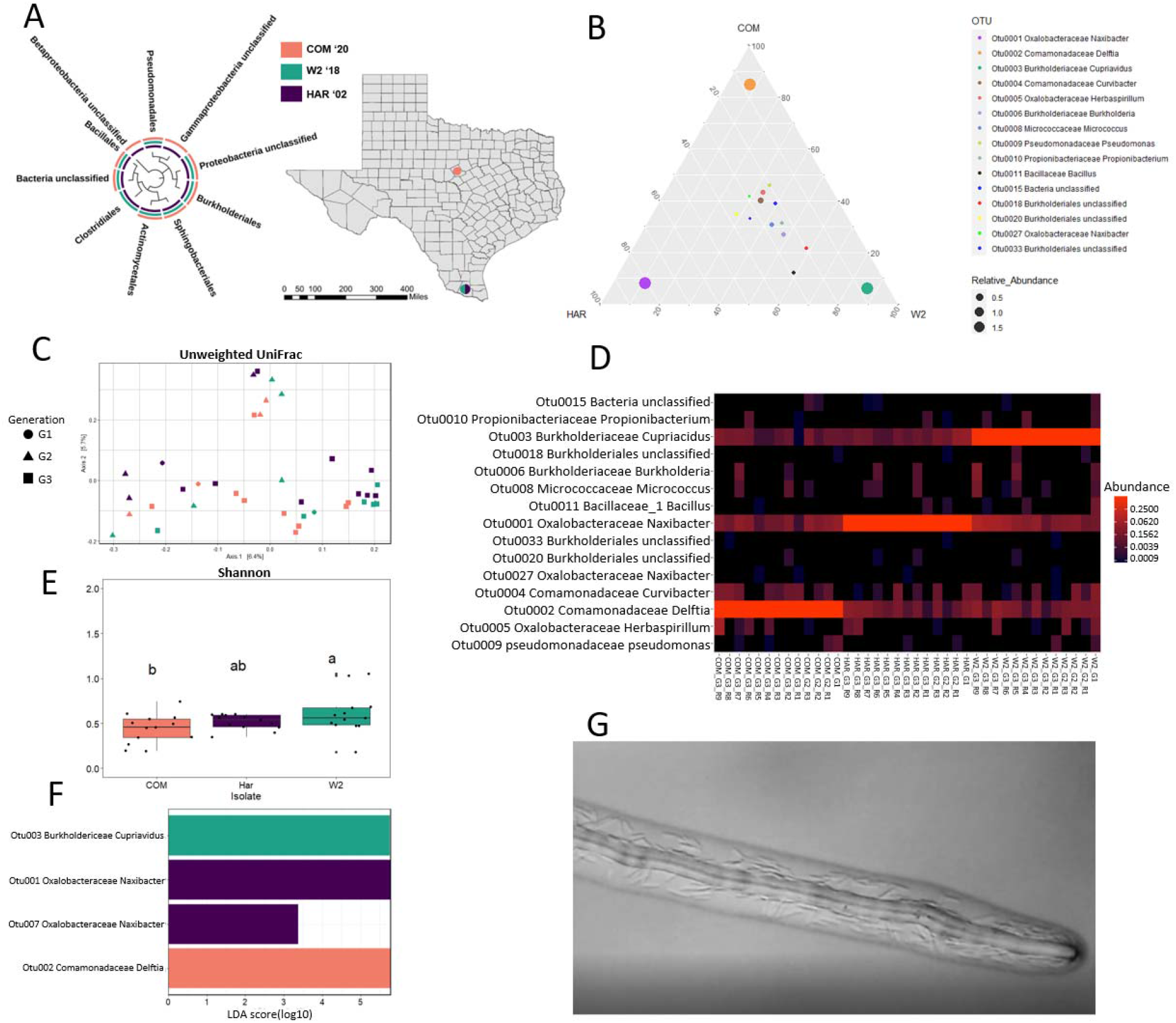

Of the ten orders that were identified, *Pseudomonadles*, *unclassified proteobacteria,* unclassified bacteria*, Actinomycetales, Sphingobacteriales,* and *Burkholderiales* were found within our three FON2 hyphosphere (Fig. 1A). The core microbiota of the FON2 hyphosphere resulted in a total of 15 OTUs, which made up 96% of the total number of OTUs. A ternary plot was used to visualize the 15 OTUs structure among the three FON2 hosts (Figure 1B). The result from the ternary plot displayed Otu004 (*Comamonadaceae*), Otu005 (*Oxalobacteraceae*), Otu006 (*Burkholderiaceae*), Out008 (*Micrococcaceae*), Otu009 (Pseudomonadaceae), Otu010 (*Propionibacteriaceae*), Otu0015 (Bacteria unclassified), Otu0020 (*Burkholderiales*), Otu0027 (*Oxalobacteraceae*), and Otu033 (*Burkholderiales*) were clustered in the center, suggesting they are strongly correlated to being a part of the core microbiome of FON2 hyphosphere regardless of spatial and temporal differences. Whereas Otu0018 (*Burkholderiales*) and Otu0011 (*Bacillaceae*) have a relatively closer relationship with W2 host compared to COM and HAR. Uniquely, Otu001, Otu002 and Otu003 which are the most abundant OTUs and found within all the samples have a strong relationship with one host.

This strong relationship of a single OTU with the FON2 host is visualized within the heatmap and shown as a high relative abundance value, further validating results observed from the ternary plot. The heatmap displays more details of Otu001 (*Oxalobacteraceae*), Otu002 (*Comamonadaceae*), and Otu003 (*Burkholderiaceae*) presence within all samples and those of the other core OTUs. The LDA reflects a similar pattern, as Otu001 has a significant relationship with HAR, Otu002 with COM, and Otu003 with W2. Further LDA results of Otu0021 and Otu007 indicate a significant relationship with HAR.

Individual FON2 isolate’s microbiome was analyzed through co-occurrence networks to compare possible host-specific assemblies (Figure 2). Within the COM network were 215 edges (33 negative and 182 positive edges), 32 nodes, HAR (1 negative and 40 positives), and W2 732 edges (4 negative and 728 positives), and 70 nodes. These assemblies’ few negative correlations were of Otu001, Otu002, and Otu003 apart from HAR, which network only included Otu001 and Otu002. Within COM, some clusters may represent a microbial assembly not correlated to the over-abundant Otu002; this similar trend is observed within W2 and HAR. Core OTU of individual isolates were identified to analyze host-specific assemblies further, these criteria identified 8 Otus within COM, 17 within HAR, and 10 with W2. The correlations among the individual FON2 core microbiota were compared, as shown in Figure 3. Similar to results from the co-occurrence model with Spearman correlation there is a negative interaction among Otu001, Otu002, and Otu003. COM dominant Otu002 is negatively correlated to the other 6 Otus. W2 dominant Otu003 negatively correlates to 7 Otus and 1 positive (Otu0019). HAR dominant Otu001 has a negative correlation with 8 other Otus and a positive correlation with 4 OTUs (Otu0014, Otu007, Otu0026, and Otu0024. Within HAR there is a positive cluster of Otu0025, Otu008, Otu0021, and Otu0013 that is negatively correlated to the dominant Otu001; therefore, may be a different assembly within the hyphosphere. Although there is a robust negative correlation among the dominant OTUs from our correlation plot, most of the co-occurrence network was positively correlated, displaying a distinct community structure.

**Fig. 2.**
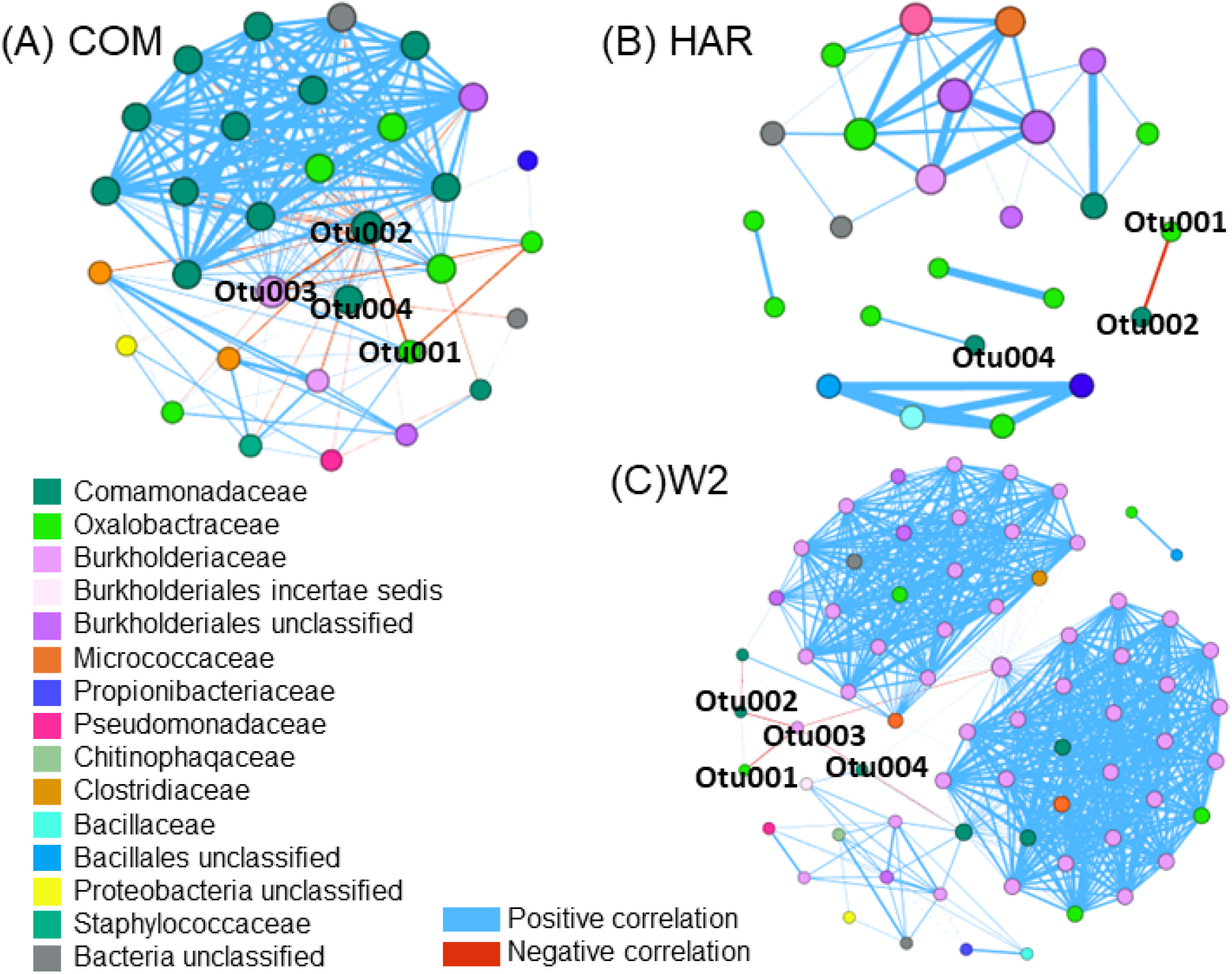

**Fig. 3.**
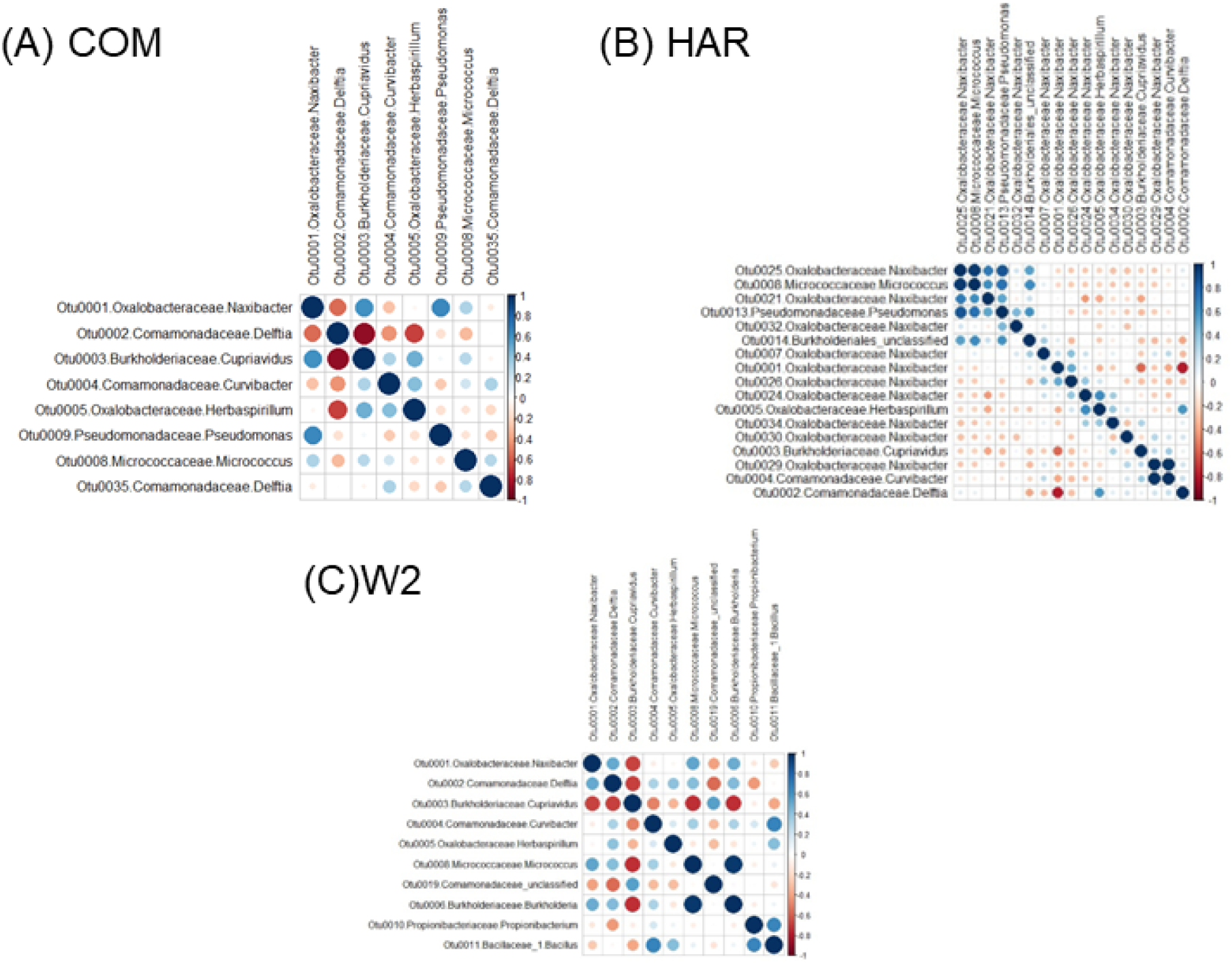

### PICRUSt results

PICRUSt analysis from 16S rRNA sequencing referenced to the KEGG database resulted 5,380 predicted KEGG Orthology (KO) groups. Of these KOs 779 were unique to COM, 150 to HAR, 69 unique to W2 and 4,137 were shared among all three as seen in figure 4. The predicted categories of major functions within the hyphosphere included Metabolism (COM at 44.99%, W2 at 48.22% and HAR at 42.71%), Environmental information processing (COM at 18.77%, W2 at 16.24% and HAR at 16.37%) and Unclassified (COM at 13.73%, W2 at 14.04% and HAR at 14.70%). In level 2 of predicted function, we chose to visualize the top 24 to compare among the three isolates hyphosphere. Within COM hyphosphere all 24 were significantly lower than HAR and W2. FON2 isolate HAR hyphosphere was found to be significantly higher to W2 in 8 predicted functions (Replication and Repair, Energy Metabolism, Cellular Process and Signaling, Enzyme Families, Genetic Information Processing, Glycan Biosynthesis and Metabolism, Biosynthesis of other secondary metabolites and Neurogenerative diseases). Within W2, four functional genes were significantly higher than those found in HAR (Amino Acid Metabolism, Lipid Metabolism, Xenobiotics Biodegradation and Metabolism and Metabolism of Terpenoids and Polyketides. Overall, these results indicate within the hyphosphere of COM (the most recent isolate) is predicted to have less shared activity compared to HAR (the oldest isolate) and W2 but has more unique unshared functions.

**Fig. 4.**
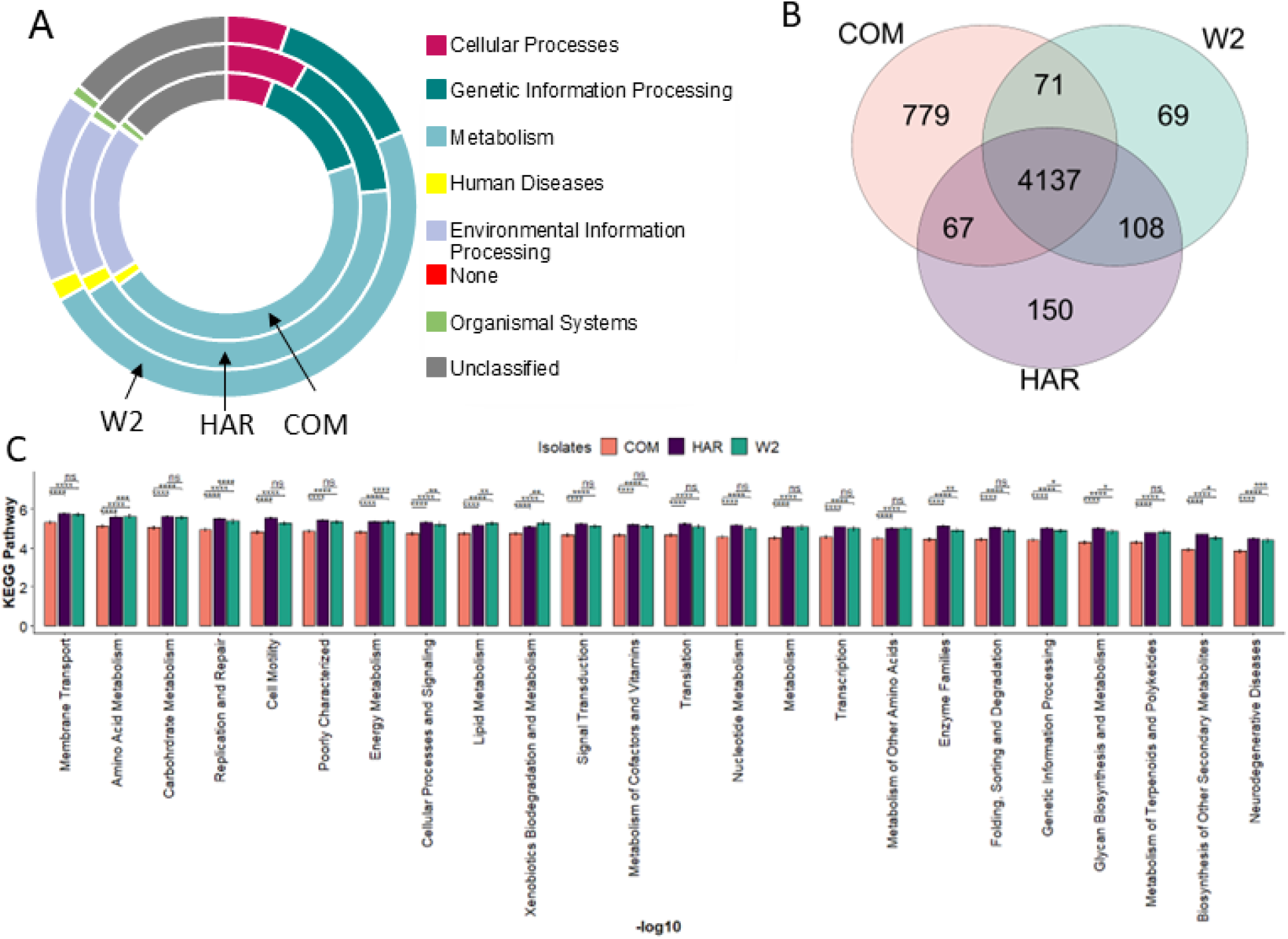

### Microscopic results

The microscope image data observed bacteria growing around the hyphae as it continued to elongate. As seen in figure 5, there is a diversity of different bacterial shapes following the elongation of hyphal growth.

**Fig. 5.**
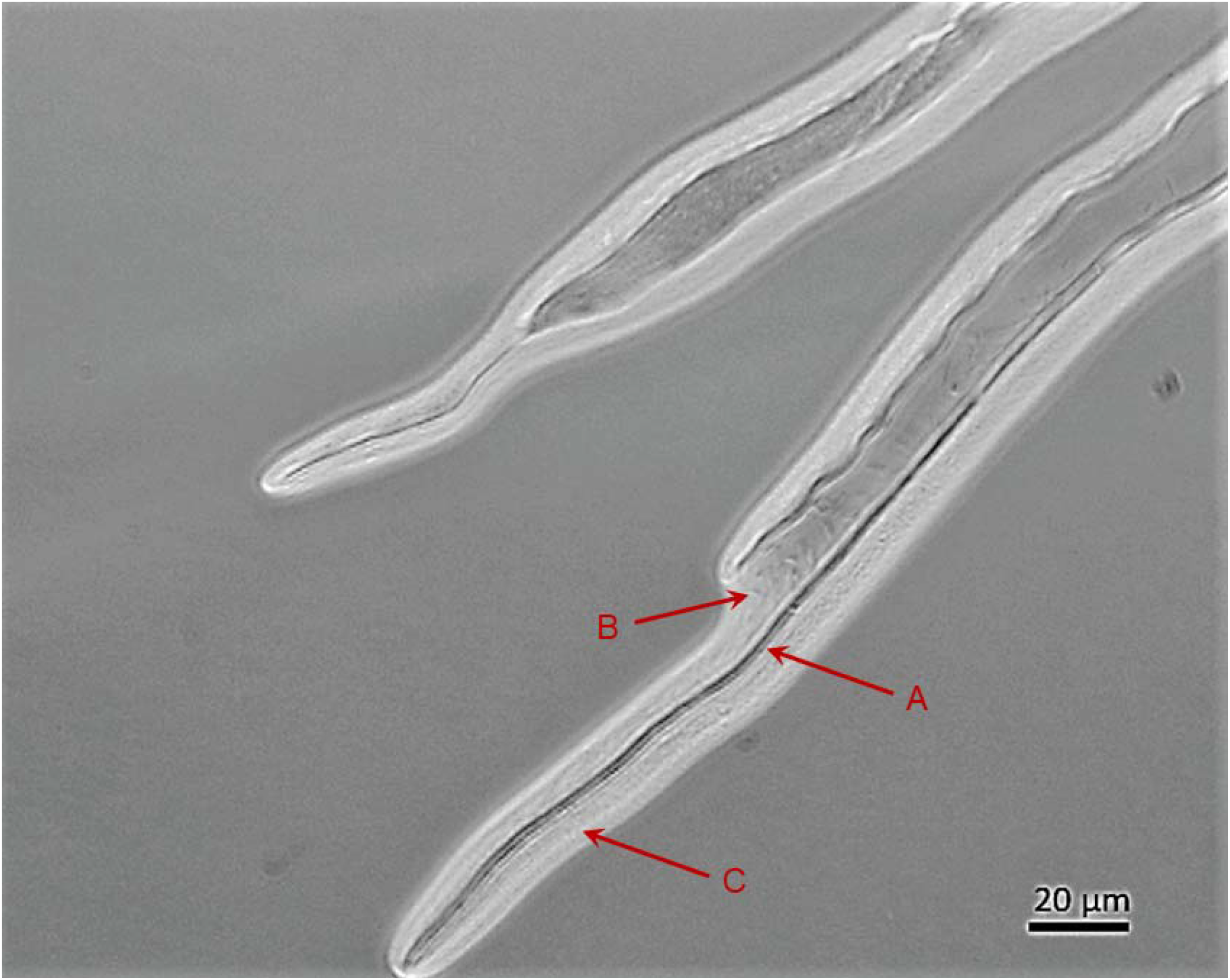
Light-microscopic images of Bacteria (B and C) surrounding the FON2 hypae (A). The acteria “C” bacteria are smaller and compact compared to bacteria “B” which are freely motile though both are moving in the direction of hyphal growth.

## Discussion

Within this study, we aimed to identify a core hyphosphere microbiota characterizing the pathobiome of FON2. Therefore, we tested our hypothesis with three isolates of different spatial and temporal backgrounds. Our three FON2 Isolates (COM from 2020, W2 from 2018, and HAR from 2016) inoculum of oatmeal and sand medium was added directly into the sterilized soil near the rhizosphere of watermelon. Unlike root dipping, this would cause less disturbance to the root and microbial system and allow for FON2 recruitment, similar to how it may recruit within the soil and host. Once symptoms were visible, stem tissue was collected to re-culture FON2 and candidate pathobiome. From the infected tissue, hyphae and bacteria would grow, and microbiota associated with the hyphal tip were re-cultured as this area specifically is the point of entrance in root infection. Results from our heatmap support that a core microbiota is associated with the hyphosphere of FON2 isolates regardless of repetition from re-culturing and differences in isolate’s backgrounds. Further results from the ternary map visualize a similar abundance of central core hyphosphere microbiota associated with the three isolates except for one dominant OTU strongly associated with a particular isolate. Thus, we have a candidate pathobiome strongly associated with the infection point which may assist in pathogen success or potentially reduce fungal antagonism from the plant and suppressive microbiota.

Our results found majority of the core microbiota was composed of the order *Burkholderiales* (i.e OTU001 *Oxalobacteraceae Naxibacter*, OTU002 *Comamndaceae Delftia* and OTU003 *Burkholderia Cupriavidus)*. Previous studies of isolates within the order include *Burkholderia terrae* BS001 which has been well studied for interactions within the hyphosphere of *Lyophyllum* sp. strain Karsten and was observed for moving along with hyphal growth but also development of biofilm around the hyphae [5]. Later, *Burkholderia terrae* BS001 was studied for the use of type three secretion system (T3SS) to be utilized in adhering to the hyphae, further results included an exchange in nutrients and protection against antifungal such as cycloheximide [38, 39]. Research specifically to isolate *burkholderia terrae* BS001 has been researched to move along the hyphae of two *Fusarium oxysporum* isolates (Fo47 and Foln3). Other pathobiome hyphae associated bacteria studied include organisms of *Oxalobacteraceae* who although has been studied to associate with arbuscular mycorrhizae, has also been observed to colonize *the* hyphae of *pythium* [40, 41]. Therefore, we suggest from our results and previous research support *Oxalobacteraceae Naxibacter*, Comamndaceae Delftia and *Burkholderia Cupriavidus* as hyphal associated bacteria of FON2. While we did not study the potential protection attributed by the hyphosphere bacteria on the FON2 strains, paralells could be drawn from literature that can suggest such potential environmental advantage provided by the bacteria. Such functions remain to be examined in our future studies.

The co-occurrence network analysis visualized a negative correlation with the dominant OTUs from each isolate and was reconfirmed through correlation plot. Although this may be due to an overabundance of these OTUs in comparison to the rest of the microbiota this may be due to a positive correlation to the fusaria that we did not test. Other than the most abundant OTUs (OTU001, OTU002 and OTU003), was OTU004 (*Comamonadaceae Curvibacter*) which as seen in fig.1D is one of the most abundant OTUs and is present in a majority of our samples. This particular OTU in the network analysis fig. 2 was near the orbit of the dominant OTUs and therefore, there may be a primary group acting as the leading pathobiome community and a secondary group that is less abundant but would take over the primary group’s functions, also known as functional redundancy. Functional redundancy is when species observed perform similar roles or functions within in community and ecosystem and therefore are substitutable to keep the ecosystem process intact [42, 43]. In summary, to maintain the pathobiome function within the hyphosphere we propose OTU001, OTU002 and OTU003 is the primary group and a secondary group including OTU004. This furthermore promotes the hyphosphere pathobiome is a functioning community and therefore warrants more research.

The rarer core microbiota of importance to consider is OTU0008 *Microccaceae micrococcus*, which was found within all three isolates hyphosphere. In previous research, members of Microccaceae have been surveyed in the rhizosphere of wheat and its potential function within the core microbiota with transferable antibiotic-resistant genes with endophytes [44, 45]. This is of interest as *Microccaceae* was identified as a core microbiota of fertilized soils in wheat systems, a studied intercrop system with watermelon for pathogen resistance [46] . In other research, *Micrococcaceae* was researched to be associated with susceptible wheat to plant pathogen *Tilletia controversa* and *Fusarium* [47]. Therefore, previous research supports our results of *Microccaceae micrococcus* is associated with the pathogen susceptible microbiota and has an association with the pathobiome of FON2 and antimicrobial assistance as was researched with *Burkholderia* should be studied. Furthermore, from our results there is sufficient evidence that *Microccaceae micrococcus* is a bacterium with fungal association.

Having established the core bacteriome associated with the hyphal tip of FON2 as the pathobiome demonstrates the complex interactions of fungi and bacteria within a pathogen system and may influence disease processes and success. Our study promotes the pathobiome concept, as strides in technology and the field of the microbiome is growing previous studies are transitioning away from the lone invader concept [48]. Furthermore research in resistant soil microbiome have proven the pathogen does not infect just the host but also infects a susceptible host microbial community and modifies it to become the pathobiome [49]. In parallel the pathogen is not infecting the host by itself but through recruitment strategies from the hosts established microbiome to form a pathobiome [50].

## Conclusion

Within this study, we characterize the core microbiota associated with the hyphal tip of plant pathogen FON2. Closely related members of our core microbiota groups have been previously researched as hyphal-associated and now present new members of *Burkholderia, Microccaceae, Bacili,* and *Pseudomonas* to be associated with hyphal interactions. As this specialized niche is the point of entry for root infection, this microbiota may contribute to essential functions of the pathobiome and may assist in disease success. Future studies that investigate the role of pathogen associated microbiota associated in host infection are required. Results from our study emphasize the importance of the pathobiome paradigm, that pathogens are no longer characterized as lone invaders but instead recruit and manipulate the hosts’ established microbiota to form a pathobiome.

### Data availability

- Sequences on SAR

## Authors contribution

The research plan for this project was constructed based on several conversations led by Sanjay Antony-Babu (SAB) with Vanessa Thomas. SAB contributed as a mentor within the project and editing of the manuscript. Vanessa Thomas carried out the experiments, analyzed the data and wrote the manuscript.

## Supplementary material

**S.1.**
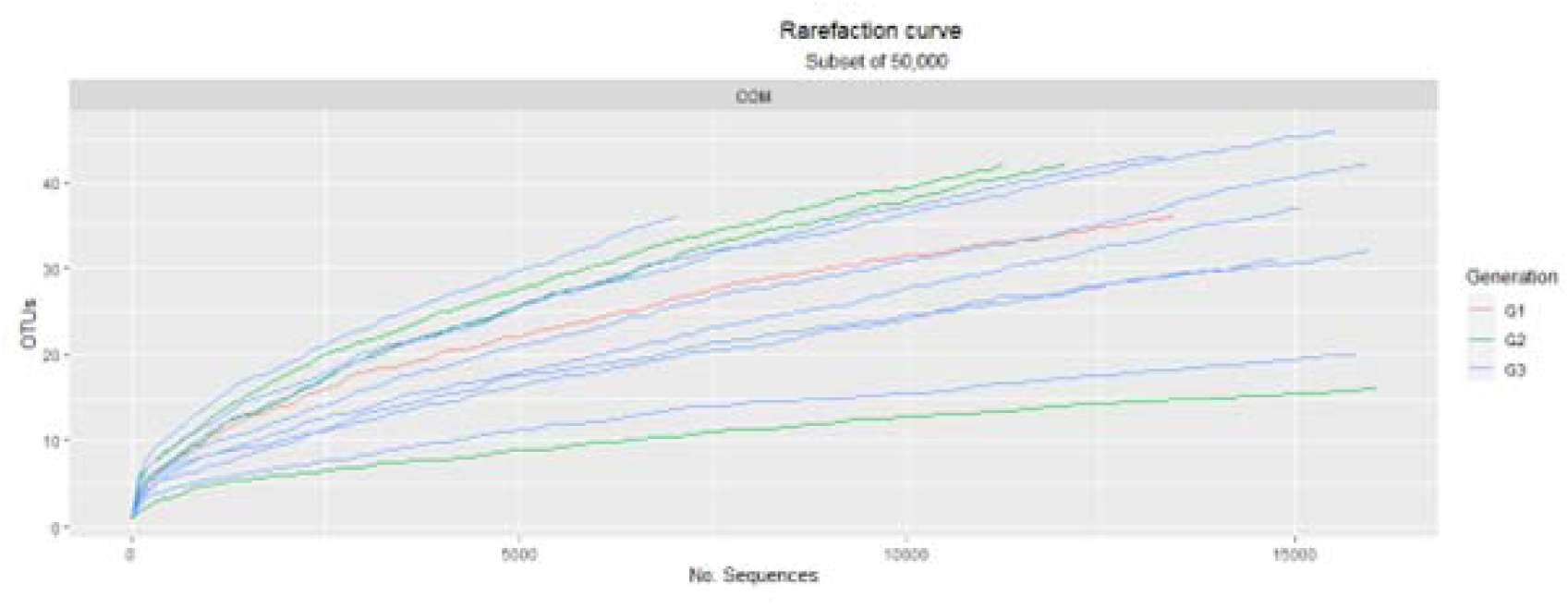

**S.2.**
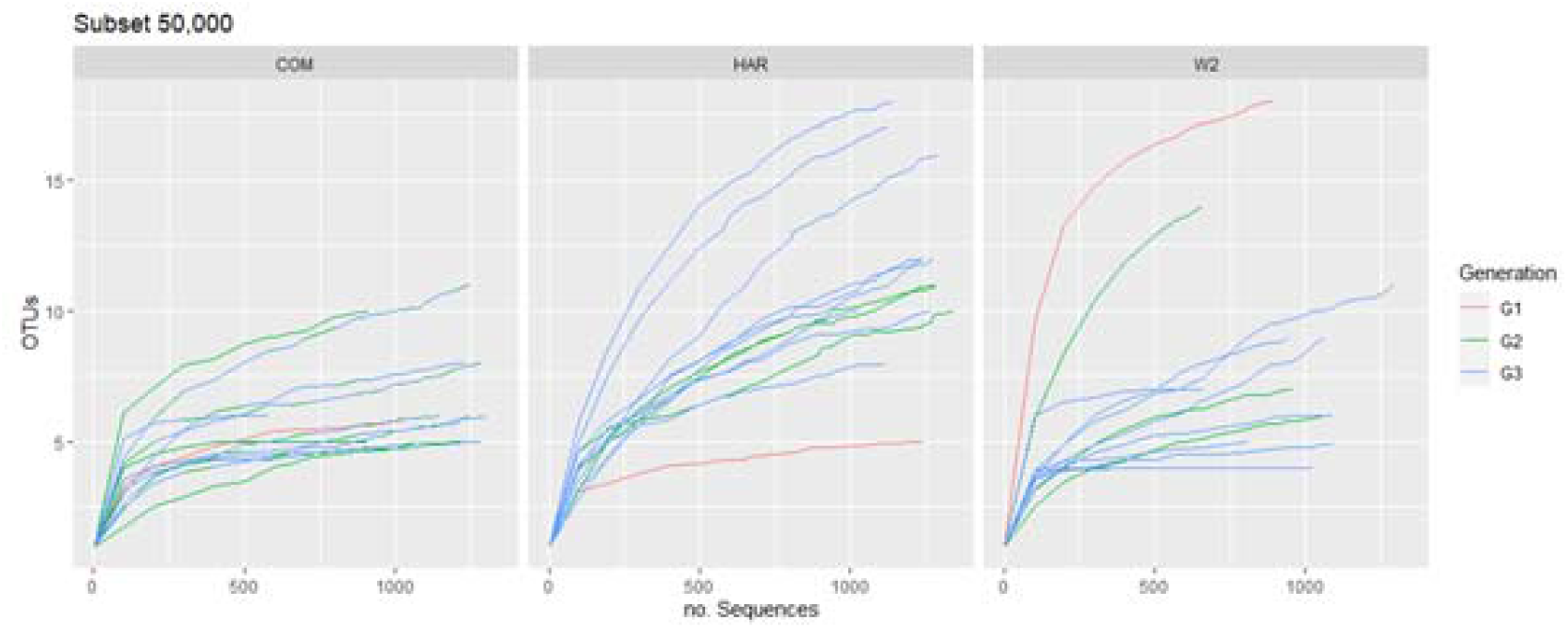

**S.3.**
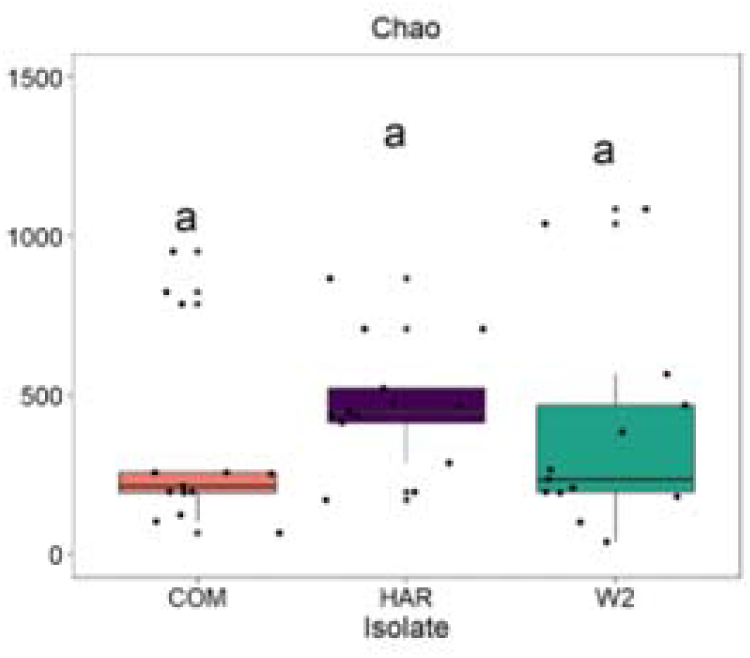

**S.4.**
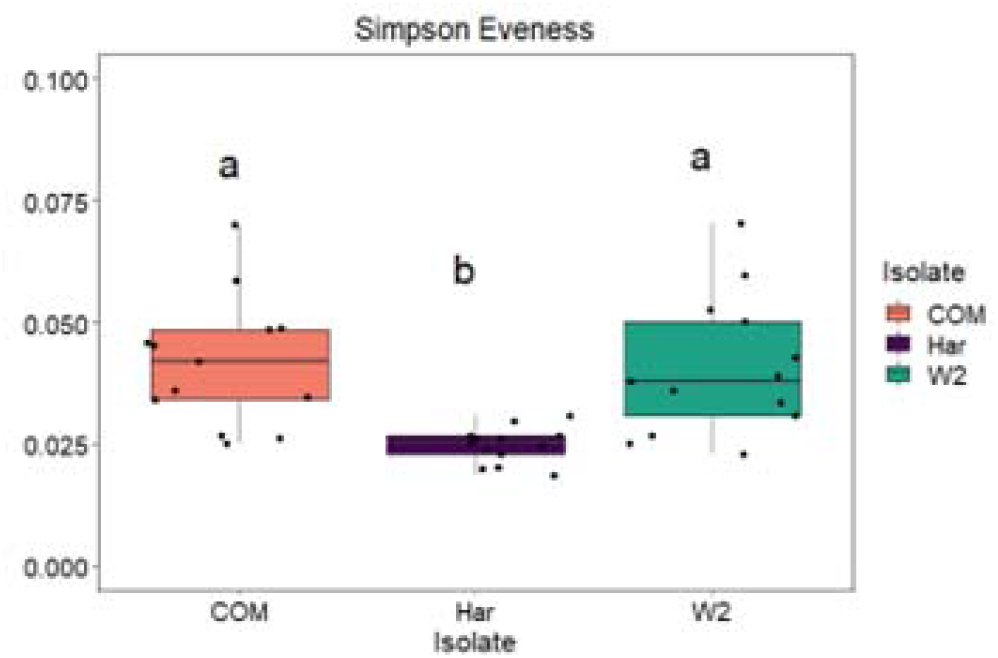

